# Modeling of Glucosinolate Biosynthesis During Biotic Stress as a Function of mRNA

**DOI:** 10.64898/2026.05.29.728632

**Authors:** Jordan Earle, Anna C.M. Neefjes, Xenja Ploeger, Marlene van Laar, Saskia C.M. Van Wees, Robert C. Schuurink, Aalt Dirk Jan van Dijk, Petra Bleeker, Huub Hoefsloot

**Author notes:** These authors contributed equally to this work (Jordan Earle, Anna C.M. Neefjes). These authors also contributed equally to this work (Xenja Ploeger, Marlene van Laar).

## Abstract

Glucosinolates are an important group of specialized metabolites in the Brassicaceae family, playing a role as defensive compounds against biotic attackers. In response to biotic stress, plants upregulate glucosinolate biosynthesis in part by increasing the abundance of enzymes in the glucosinolate biosynthetic pathway. As an increase in enzyme abundance is often preceded by an increase in the corresponding mRNA levels, the dynamic changes in mRNA levels should capture the information required to infer how metabolite levels change over time. In order to test this hypothesis, a time series of experimental glucosinolate content data collected from *Arabidopsis thaliana*, exposed to either a mock or methyl jasmonate (MeJA) treatment, as a proxy for biotic stress, was combined with existing mRNA abundance data over time at the same developmental stage and treatment. We propose the GEEM model, a multilevel mechanistic ordinary differential equation (ODE) model, which goes from Gene expression to an enzyme level model, followed by a Michaelis Menten kinetics metabolite model, to simulate the dynamics of a segment of the indolic glucosinolate pathway. In order to constrain the GEEM model, three models were fit to experimental *de novo* specialized metabolite data, using different degrees of freedom by utilizing both a Gradient Boosted Tree model with a tested architecture to predict the kinetic constants, and augmenting these predictions with a literature review of the known Michaelis Menten kinetic constants from the glucosinolate pathway. Using Sequential Monte Carlo - Approximate Bayesian Computing to fit the GEEM model to the experimental data, we showed that given the mRNA levels and initial concentrations of metabolites, the changes in specialized metabolites over time and treatment can be modeled.

**Author Summary:** We study how plants adjust their natural chemical defenses over time when they are under attack from living organisms. In the mustard family, including the subject of our experiment Arabidopsis, one important group of defense chemicals is called glucosinolates. When Arabidopsis is under attack, certain gene pathways can be activated or deactivated, allowing the plant to modulate the amount of enzymes they produce, which in turn modulates the levels of these defensive chemicals.

In this work, we combine measurements of gene activity and glucosinolate levels from Arabidopsis treated with a compound used in stress signal that mimics insect or pathogen attack. We then constructed a mathematical model that goes from gene activity, to amount of enzyme present, and ends with the amounts of specific glucosinolates over time.

By fitting this model to experimental data, we show that it is possible to predict how glucosinolate levels change over time from the gene activity and initial glucosinolate levels. Our approach offers a way to connect gene expression datasets to real changes in plant defense chemistry, with potential applications in plant breeding and insight into how these pathways change due to stress.

## Introduction

Understanding the interplay between the genome, environment and a plant’s phenotypical characteristics is of key interest in biological sciences, and is relevant when considering ever changing environmental conditions. In order to defend themselves, plants can produce specialized metabolites, which can cause a number of effects on the herbivores and pathogens. The concentrations of these specialized metabolites can vary under different environmental and stress conditions due to alternate prioritization of resource allocation. The ability of plants to fine-tune the production of specialized metabolites is crucial for effective defense responses and environmental adaptation. A well studied group of specialized metabolites are glucosinolates (GSLs), which are exclusively present in the Brassicales order, containing crops species like broccoli, cabbage, radish and brussels sprouts, but also the model species *Arabidopsis thaliana* [1]. GLSs typically share a common structure comprising a *β*-thioglucose moiety, a sulfonated oxime moiety, and a variable side chain and can be classified into three major groups based the amino acid precursor of the variable side chain [2]. Indolic GLSs are derived from tryptophan, aliphatic GLSs from methionine and aromatic GLSs from phenylalanin or tyrosine. In Arabidopsis the indolic group consist of five different GLSs, Indole-3-ylmethyl Glucosinolate (I3M), 1-hydroxyindole-3-ylmethyl Glucosinolate (1OHI3M), 4-hydroxyindole-3-ylmethyl Glucosinolate (4OHI3M), 1-Methoxy-3-indolylmethyl Glucosinolate (1MOI3M), 4-Methoxy-3-indolylmethyl Glucosinolate (4MOI3M), which will be of focus in this work (Fig 1).

**Fig 1.**
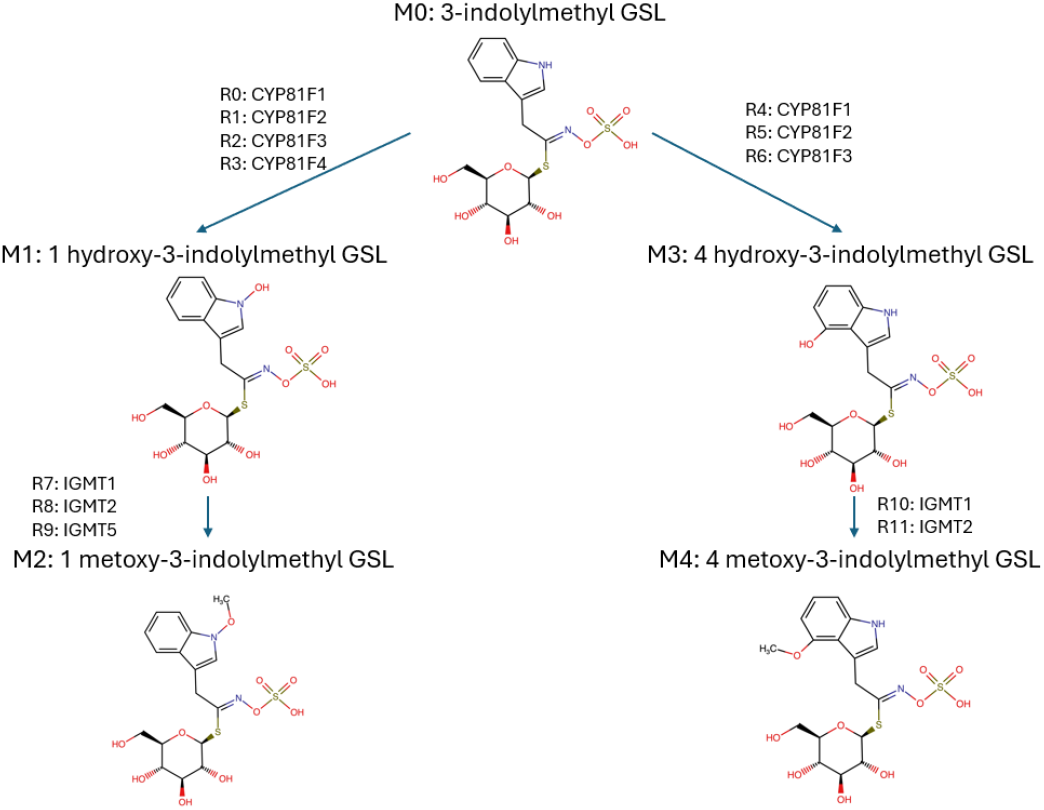
Metabolic pathway of the modeled indolic glucosinolates. Figure shows the GSLS modeled: Indole-3-ylmethyl Glucosinolate (I3M), 1-hydroxyindole-3-ylmethyl Glucosinolate (1OHI3M), 4-hydroxyindole-3-ylmethyl Glucosinolate (4OHI3M), 1-Methoxy-3-indolylmethyl Glucosinolate (1MOI3M), 4-Methoxy-3-indolylmethyl Glucosinolate (4MOI3M)) and the various enzyme/reaction combinations modeled (CYP81F1-4, IGMT1, IGMT2, IGMT5). Each metabolite and enzyme/reaction combination is represented in a shorthand (M# and R# respectively).

Upon insect herbivory, GLSs are hydrolysed by the enzyme myrosinase, triggering a so called “mustard bomb”, and releasing toxic products that act as insect deterrents. Plants can modulate their GLS profiles in response to different biotic stressors [3–7], sometimes accomplished through multiple enzymatic reactions for the same substrate to product, but the mechanisms by which these profile shifts are regulated remain poorly understood. It has been demonstrated in Arabidopsis that indolic glucosinolates and the genes encoding enzymes in the glucosinolate pathway increase when exposed to MeJA, which is converted in the plant into Jasmonic Acid (JA) [3–6]. JA is a key phytohormone that is increased in plants upon feeding by herbivorous insects and causes major shifts in gene expression and metabolite production [8]. Mechanistic models can provide insight into the choices plants make upon receiving a stressor, giving insight into the altered fluxes and metabolite profiles within the indolic GLSs pathway.

To understand the characteristics of enzyme-metabolite reactions, mechanistic models using mass action kinetics or Michaelis Menten kinetics have been used to characterize the rates of metabolites synthesis in biosynthesis systems. These kinetic constants are often found experimentally, but this is not always feasible and can be costly, due to the time and resources required to produce the required enzymes and substrates for the reactions. In these situations, the same mechanistic models which were previously used to understand the dynamics of the enzyme substrate reactions can be used to determine the missing kinetic constants via fitting to easier to obtain experimental data rather than performing traditional enzyme kinetics experiments. These mechanistic models can also be used in conjunction with traditional enzymatic assay analysis in order to reduce the amount of assays required to determine the kinetic constants [9].

Many enzyme kinetic models assume constant enzyme concentrations, thereby simplifying the systems so the kinetic parameters become the only dependent variable of metabolic flux [10, 11]. In vivo, however, enzyme concentrations are highly dynamic, organisms continuously adjust their metabolic content in response to developmental cues and biotic and abiotic conditions. The biosynthesis of specialized metabolites is typically low under non-stress conditions and is transcriptionally activated when required, for example upon herbivore attack or pathogen infection [12]. Upon perception of a stressors a signaling cascade (often involving phythormones such as jasmonic acid or salicylic acid) is triggered that changes gene expressions leading to increased production of biosynthetic enzymes leading to elevated specialized defense-related metabolites. As such, a model could capture the changes in glucosinolate content via the observed changes in mRNA levels of the biosynthetic enzymes by incorporating variable enzyme levels, given the initial mRNA, enzyme and metabolite concentrations, and the mRNA concentration through time.

Experimentally determining metabolite dynamics under stress is labor intensive. Computational models that integrate gene expressions, enzyme kinetics, and metabolite dynamics can be used to investigate the impact of specific stressors and guide more targeted experiments. By understanding how enzyme levels and specialized metabolite content can change over time, such models can improve our understanding of resource allocation and the structure and control of biosynthesis networks during stress response. For instance, in the work of Cloutier et al. [13], the metabolic network of Arabidopsis was modeled through time using mRNA levels to show how single gene perturbations affected seed fatty acid content. However, these models did not focus on how different (combinatorial) environmental conditions would affect the fatty acid profile.

We propose that the changes in gene expression during an (a)biotic stress event are reflected in the changes of metabolite concentration. As existing models tend to used a fixed enzyme concentration to find kinetic constants. Changes in enzyme levels are most often preceded by changes in the corresponding mRNA levels. Therefore, to capture dynamics during a stress event, changes in gene expression and the resulting changes in enzyme concentrations, must be considered. By considering the enzyme concentration as a function of the corresponding gene expression, enzyme kinetics models should be able to capture the up and down regulation of enzymes and metabolites through time over different conditions.

In order to test this hypothesis, we created a model which uses de-novo metabolite data consisting of four indolic GSLs, collected through time from rosette tissue of Arabidopsis (Col-0) combined with an existing time series of mRNA data [14], both from Arabidopsis treated with MeJA. This Gene Expression, Enzyme, Metabolite (GEEM) model combines mass action kinetics and Michaelis Menten kinetics to replicate experimental observations from the segment of indolic glucosinolate pathway using the gene expression data. The mRNA levels are used to model the production of the enzymes using mass action kinetics. By having the enzymes modeled such, we do not require the assumption that one mRNA count is equivalent to one protein, allowing the data to constrain the relationship between the mRNA and enzymes. These variable enzyme levels are then used with combinations of Michaelis Menten and mass action kinetics to model the changes in metabolite concentration through time.

The GEEM model requires kinetic constants for every step in the process for every enzyme and substrate. As there are no experimentally obtained kinetic constants for the enzyme-substrate reactions in question, the architecture from recent machine learning models from Kroll et al. [15, 16] was utilized to predict the Michaelis constant *K*_*M*_ and turnover rate *k*_*cat*_, as a starting point. This gradient-boosted tree model was trained on a subset of the original dataset containing specialized metabolite reactions of spermatophythes (seed plants).

After evaluating the ability of the GEEM model to reproduce the experimental data using the machine learning model predicted reaction kinetics and fitting the enzyme mass-action parameters to the data, we subsequently fit the Michaelis–Menten constants with the enzyme mass-action parameters, constraining them around the machine learning model predicted values. Next, a literature study was performed on known Michaelis Menten kinetics of the enzymes involved in the production of glucosinolates in Arabidopsis. This information was combined with the previous machine learning model prediction based priors and the model was fit for comparison. The models proved able to replicate the experimental metabolite content results given the precursor metabolite profile and treatment specific mRNA level profiles.

## Materials

To test whether changes in gene expression under (a)biotic stress are mirrored by changes in metabolite concentrations, we generated a targeted metabolomics dataset under growth conditions and a sampling protocol closely matching those used to generate the transcriptomics data [14]. *Arabidopsis thaliana* of the ecotype Columbia-0 (Col-0) were grown as described in the supplemental methods. In short, five week old plants were treated with either a MeJA (0.1 mM) or a mock solution (see supplemental Metabolite Data Collection). Leaf material (n=6) was sampled and analyzed by targeted LC-MS for GLS content.

In the *de novo* metabolite experiment, six replicates were sampled at 2, 4, 8, 24, 48, 120 and 216 h after treatment, as well as immediately after treatment (0 h) and 1 h prior to treatment (-1 h). For each replicate the rosette was harvested and flash frozen in liquid nitrogen. The GSLs of interest (I3M, 4OHI3M, 1MOI3M, and 4MOI3M) were extracted and the concentration measured using the procedure outlined in the supplementals. A standard for 1OHI3M was not available and it has not been observed in measurable quantities in literature [17, 18]. The resulting data for these metabolites can be seen in Fig 2.

**Fig 2.**
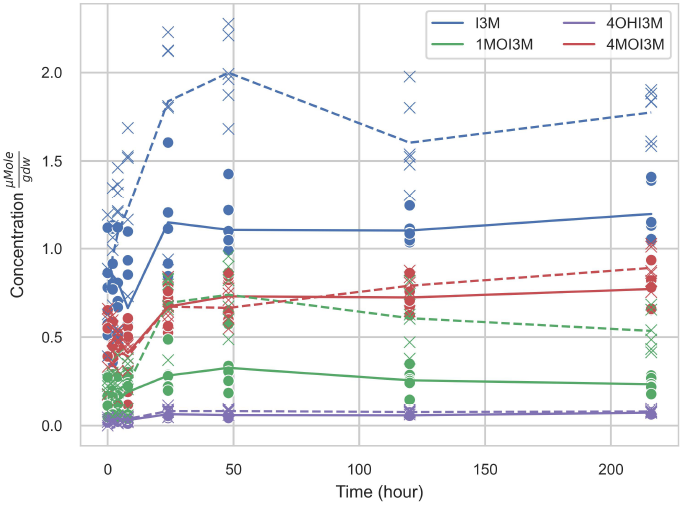
Experimentally obtained glucosinolate content. 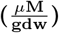 Figure shows the mean concentrations of the glucosinolates, I3M, 1OHI3M, 1MOI3M, and 4MOI3M, over time after the mock (solid lines) and MeJA (dashed lines) treatments. Individual replicates for the mock (circles) and the MeJA (crosses) are shown (n=6).

The transcriptomic data used in this experiment comes from Hickman et al. [14], where a similar procedure was followed for the growth and treatments. The normalized mRNA counts were determined for the genes encoding the particular enzymes of interest (Fig 1 CYP81F1-F4, IGMT1, IGMT2, IGMT5) as described in Hickman et al. [14]. In this mRNA level dataset, the 6th leaf was sampled 0, 0.25, 0.5, 1, 1.5, 2, 3, 4, 5, 6, 7, 8, 10, 12, 16 h after treatment. The mRNA level data for both treatments can be seen in Fig 3.

**Fig 3.**
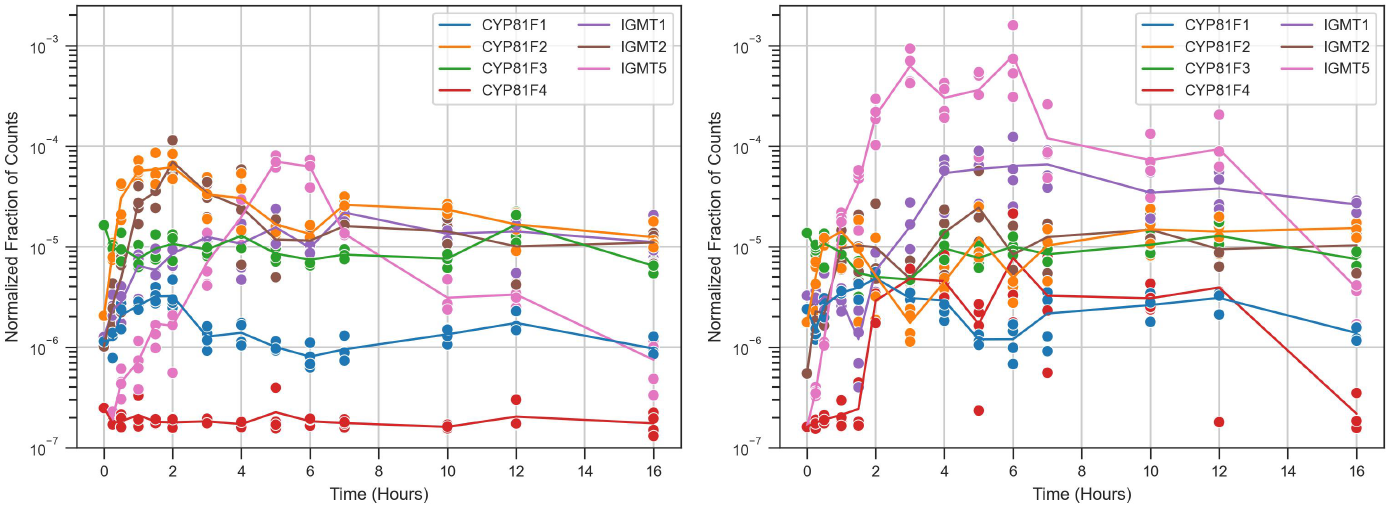
Transcript abundance enzymes of interest after Mock and MeJA Treatment. Figures show the experimental observations of the normalized fraction of counts (log scale) for the enzymes of interest in the mock (a/left) and MeJA (b/right) treatment. Lines show the mean mRNA levels across time; individual data points are shown as symbols.

## Methods

### ML Kinetic Constants Predictions

To begin to constrain the undefined Michaelis Menten kinetics, two XGBoost tree models with architectures derived from Kroll et al. [15, 16] were used to predict the *K*_*M*_ and *k*_*cat*_ of the enzymes of interest. An analysis of the original *k*_*cat*_ model showed that it was unable to predict meaningful values for mutants and unfamiliar enzymes [19]. Therefore, we retrained the architecture using a subset of the original data, focused on the specialized metabolites biosynthesis from seed plants, in order to bring the training data closer to that required for the task.

#### Model datasets

The original models from Kroll et al. [15, 16] integrated kinetic values from three different databases, i.e. Brenda, UniProtKB, and SABIO-RK, covering a broad range of organisms and reaction types in order to predict *K*_*M*_ and *k*_*cat*_. For this model we utilized SABIO-RK and UniProtKB, as Brenda contained only duplicates for the enzymes of interest. Different organisms can have different metabolic dynamic scales, and different types of reactions, such as primary, specialized, or redox reactions, can have vastly different speeds. As such, to predict the enzyme kinetics for a specific functional type of reaction and organism, the training data from the original model contains information which can skew the prediction. Here, we are only interested in a subset of enzymes involved in part of the indolic glucosinolate (GLS) biosynthetic pathway of *Arabidopsis thaliana*. Thus, we trained a new predictive model using a mixture of architectures from Kroll et al. [15, 16], focused on specialized metabolism enzymes from seed plants, bringing the training data closer to the specific task.

#### Model Inputs

As the function of an enzyme is related to the temperature and the pH of the environment of the reaction, the new metabolite specific machine learning models use the temperature and the pH from the experimental source of the kinetics. For our predictions, we used the average pH found in the literature experiments, i.e. pH 7.5, and the temperature of the experimental conditions, i.e. 21 °C. Additionally, the model used to predict the *K*_*M*_ incorporates the main substrate, enzyme and reaction. The main substrate is incorporated by training a graphical neural network (GNN) to create a representation of the molecule structure, more information can be found in Kroll et al. [15, 16]. The enzyme representation is created using the ESM-2 [20] model and the reaction embedding is created using RDkit [21]. The model used to predict the *k*_*cat*_ uses the enzyme representation and its reaction representation, using the aforementioned methods, in addition to the temperature and the pH.

#### Model Predictions

Using the seed plant and specialized metabolite specific models, the kinetic constants for the enzyme reactions were predicted (Table 1). These values were compared to the known literature values for GSL reactions, (Table 3) and were found to lie within the observations from literature. Those comparing the tables may notice the max value of *K*_*M*_ was limited to 500 *µM*, but there have been observed values as high as 80000 *µM* [22].

**Table 1.** Machine learning models Michaelis Menten Kinetics predictions. shows the reaction number, enzyme - protein combination and machine learning models predictions per reaction for *K*_*M*_ and *k*_*cat*_.

| Reaction | Enzyme - Product | $K_M(\mu M)$ | $k_{cat}(h^{-1})$ |
| --- | --- | --- | --- |
| **R0** | CYP81F1 - 1OHI3M | 5.30 | 250.77 |
| **R1** | CYP81F2 - 1OHI3M | 6.36 | 1139.04 |
| **R2** | CYP81F3 - 1OHI3M | 6.09 | 563.13 |
| **R3** | CYP81F4 - 1OHI3M | 5.93 | 1039.98 |
| **R4** | CYP81F1 - 4OHI3M | 5.30 | 245.82 |
| **R5** | CYP81F2 - 4OHI3M | 6.36 | 928.03 |
| **R6** | CYP81F3 - 4OHI3M | 6.09 | 429.62 |
| **R7** | IGMT1 - 1MOI3M | 42.83 | 195.11 |
| **R8** | IGMT2 - 1MOI3M | 38.83 | 295.31 |
| **R9** | IGMT5 - 1MOI3M | 50.68 | 260.32 |
| **R10** | IGMT1 - 4MOI3M | 114.10 | 187.05 |
| **R11** | IGMT2 - 4MOI3M | 130.10 | 283.12 |

### Indolic Mechanistic Model

The GEEM model is a multi-system model which attempts to capture the changes in metabolite concentration as a function of the gene expression. Starting at the mRNA level, the change in the rate of synthesis of the enzymes is modeled. The change in enzyme concentration then modifies the rate of synthesis of the metabolites, and therefore the metabolite content becomes a function of the mRNA level over time. The different systems are modeled with ODE’s to reflect the rate of production and loss of the enzymes and metabolites through time.

To test the model, we are examining a segment of the indolic GLSs pathway, seen in Fig 1. The GEEM model simulates from the start of the experiment (t=0) to the last metabolite measurement (t=216h). To determine the combinations of parameters that would allow for both conditions to be modeled given the condition specific mRNA level, the GEEM model was fit simultaneously to both the mock and MeJA treatments. In order to achieve the model which could best reflect the data, the priors of the Michaelis Menten kinetic constants were modified over three experiments to include more information/freedom into the model. In the first version, the ML parameters were used as fixed values, and the remaining enzyme kinetic parameters were fit. In the second version, the ML predicted constants were used to guide the priors of the parameters. In the third and final version, the ML based priors were combined with existing literature knowledge of other enzyme reactions in the GSL pathway to create the final priors.

All code and data for the model will be available on Github.

### Enzyme Modeling

#### Enzyme Model

The rate of change of the concentration of enzymes is modeled as a function of mRNA level using mass action kinetics as proposed in Cloutier et al. [13]. The enzymes are synthesized from their associated mRNA and lost at a rate proportional to the amount of enzyme present. This scheme can be seen for the general case in eq. 1.

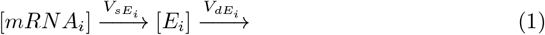

The rates equation derived from the scheme can be seen in equation 2. *V*_*sE*_ is the rate of synthesis of the enzyme as a function of the mRNA concentration while *V*_*dE*_ is the rate of loss of the enzyme which is a function of the enzyme concentration.

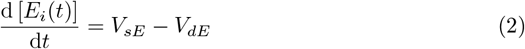

By assuming mass action, *V*_*sE*_ and *V*_*dE*_ can be re-written as seen in equation 3 and 4. The rate of synthesis (*V*_*sE*_) is then proportional to the concentration of mRNA and the synthesis reaction rate constant 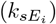 for each synthesis reaction. Likewise, the rate of loss (*V*_*dE*_) is proportional to the concentration of enzyme and the loss reaction rate constant 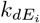 for each synthesis reaction.

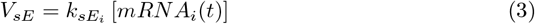

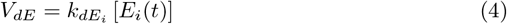

Combining, we arrive at the generalized rate equation for the enzyme kinetics (eq. 5).

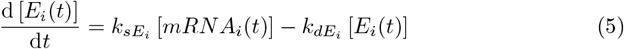

#### Model Inputs

In order to model the concentration of enzymes over time, using eq. 5, the mRNA level, represented as a concentration, through time is required as an input. The data from Hickman et al. [14] is discrete, while the model requires a continuous input. As such, a linear interpolation of the data is used to ensure a continuous data source.

The mRNA data includes measurements until 16 h. As the model simulates until t=216 h, we need to make assumptions for what occurs with the mRNA levels over that time. The mRNA data indicates that by 16 h, mRNA levels of the GSL enzymes were returning to approximately their initial values. Literature also suggests that the mRNA level response should be returning to pre-stress levels around 24 h [23]. As such, we assume that at t=24 h the mRNA levels return to their pre-stressed values and remain such for the rest of the simulation, in line with the quasi steady state assumption we made at the start of the experiment.

Additionally, for some specific mRNAs, counts close to 0 were observed. To be able to still use these data points, these low values were substituted for 10 counts, as done previously by Hickman et al. [14].

The model uses a concentration of mRNA rather than the counts found in the transcriptomics analysis. To convert the fraction of counts (*c*_*n*_) data into a concentration of mRNA, a conversion was applied as seen in eq. 6. We assume the number of mRNA molecules per cell (*mRNA*_*c*_ = 250000) taken from literature [24, 25] of mammalian cells in absence of data on plant cells. The assumed number of cells per leaf (*nc*_*l*_ = 764, 000) originates from literature [26–28] and the mass of a leaf (*m*_*l*_) was measured in the experiments and averaged to 35 *mgFW*, which can be converted to 4.697 *mgDW* using the conversion factors from Tolleter et al. [27]. *N*_*A*_ is the Avogadro constant which is used to convert the number of particles to the amount of a substance (in moles).

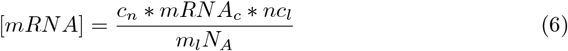

#### Additional Model Constraints

In order to determine the rate of change for each enzyme, the rate of synthesis with respect to the mRNA level and the rate of degradation of the enzymes are needed. The synthesis rate of different families of enzymes (*V*_*sE*_) [29] and the degradation constant of proteins (*k*_*dE*_) [30] in Arabidopsis were used for fitting the model (Table 2). The synthesis rate provided in Piques et al. [29] is in terms of enzymes/minute, which is the overall rate, unrelated to mRNA level. We assume that at the start of the experiment, before perturbation, the plant is in a qquasi steady state 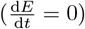. Therefore, in order to convert the synthesis rate (*V*_*sE*_) into the synthesis reaction rate constant (*k*_*sE*_) we use the concentration of mRNA at t=0 as the steady state value. Next, applying first order mass action approximation we solve for *k*_*sE*_ as seen in equation 7.

**Table 2.** Enzyme model priors and distributions. shows the reactions, prior distribution ranges, and prior distributions types used for fitting for the synthesis rate (*V*_*sE*_) and degradation constant (*k*_*dE*_) derived from literature [29] [30]

| Parameter | Reactions | Range | Prior |
| --- | --- | --- | --- |
| $V_{sE}$ | R12 - R18 | $[3.325e^{-7}, 0.1015] \mu M h^{-1}$ | log-uniform |
| $k_{dE}$ | R12 - R18 | $[0.00208, 0.02042] h^{-1}$ | uniform |

**Table 3.** **Michaelis Menten kinetics prior ranges and distributions**the reactions, prior distribution ranges, and prior distributions types used for fitting for *K*_*M*_ and *k*_*cat*_ derived from literature.

| Parameter | Reactions | Range | Prior |
| --- | --- | --- | --- |
| $K_m$ | R0 - R11 | $[1e^{-3}, 500] \mu M$ | log-uniform |
| $k_{cat}$ | R0 - R11 | $[3.6, 1.8e^5] h^{-1}$ [35] [36] | log-uniform |

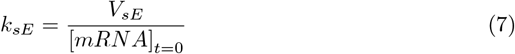

In a similar method, we solve for the initial concentration of enzyme from equation 5, using the assumption of quasi steady state, as seen in eq. 8.

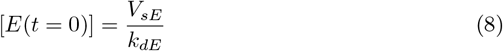

While it could be the case that some of the genes in question are linked to the circadian clock [31], in this experiment we assume these effects are negligible as there is not enough data to incorporate these effects into the model.

### Metabolite Modeling: Michaelis Menten Model

To model the metabolite synthesis rate, Michaelis Menten kinetics, a standard approach to modeling enzyme-substrate synthesis, are used. The fate of the indolic GLSs after production is not well known (e.g. whether they degrade, are recycled, transported outside of the leaf, dispersed through growth of the leaf, etc). Therefore, we model the loss of the metabolites using mass action kinetics, which could represent any combination of these fates. The system can then be expressed as a rate of synthesis 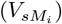 and loss 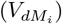 of metabolites as in equation 9.

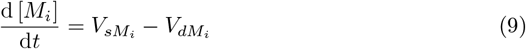

The synthesis reaction is modeled using Michaelis Menten kinetics (equation 10) with the ith metabolite being produced from the hth metabolite by the jth reaction using the turnover rate 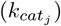 and Michaelis Menten constant 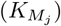 for the specific enzyme (*E*_*j*_). As there are multiple reactions, these are summed over the enzymes involved in the synthesis of the substrate (*n*_*sE*_).

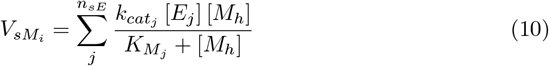

The loss rate can have two components (equation 11), the first being the synthesis of the metabolite into another product, another Michaelis Menten reaction. Here the ith metabolite is being used in the kth reaction by enzyme k. This is summed over the enzymes involved in the loss for the substrate (*n*_*lE*_). The second is the general loss of the metabolite modeled through mass action kinetics where the loss of the ith metabolite is proportional to the loss rate 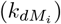 and the concentration of the metabolite.

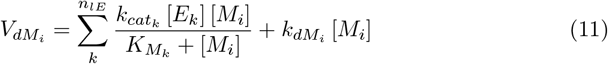

These components are then combined for the different synthesis and loss reactions for a given metabolite. In equation 12 we see an example for a reaction which has one source, one product and a loss. Here we can see the summation for the synthesis and the loss of the metabolites respectively.

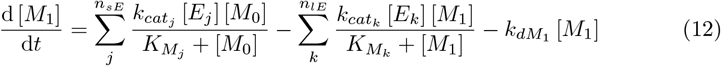

Each metabolite is controlled by one such equation, combining all the synthesis and lose reactions for the enzymes involved.

#### Model Input

To simulate metabolite dynamics over time, the model requires a precursor metabolite that acts as a deterministic input to the system. This requirement stems from the system structure. If the source is not a constant supply, or a deterministic (function based) supply without loss, the precursor can be used up completely during the experiment. Once the precursor metabolite is completely used by the reactions, the other metabolites would also start to degrade, as no more production can occur. This breaks the ability for the system to have a non-zero steady state and therefore in this experiment we make I3M the source, using the mean time trajectory at each measurement per treatment. Since the model is concerned with capturing the average system dynamics, the initial conditions of other GSLs are set to the average of each at t=0 h.

As 1OHI3M was not recorded during the analysis and has not been recorded in previous literature in Arabidopsis, an assumption about the concentration of this metabolite was made, in order to constrain the model and metabolite. The glucosinolates are thought to have similar dynamics, considering that many of the same enzymes are used for both reactions. Therefore, we make the assumption that the concentrations between similar reactions are on a similar scale as well. As such, in this model in order to determine the concentration of 1OHI3M, the ratio between the concentrations of 4OHI3M and the concentrations of 4MOI3M was calculated at t=0 h and using the concentration of 1MOI3M as seen in eq. 13 the concentration of 1OHI3M was approximated. The concentration found was 1.333*e*^*−*2^ *µmol/gDW*. This was assumed to be constant for the course of the time series, as the desire was to ensure a low concentration was maintained.

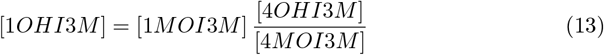

#### Additional Model Constraints

The metabolite loss rates are related to the chosen synthesis parameters, and in order to reduce the number of parameters being fit, the assumption of near homeostasis is used to determine the rates of metabolite loss. Assuming that the change is almost 0 over time at the initial conditions, we can rewrite the basic metabolic reaction equation (eq. 12) to solve for the rate of metabolite loss 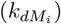 for each metabolite as seen in equation 15.

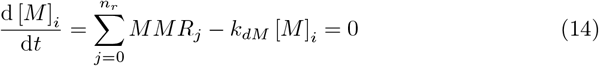

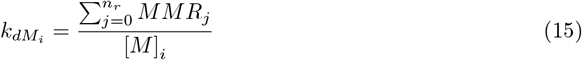

Each metabolite has a number of reactions (*n*_*r*_) where the metabolite is either synthesized or converted into another metabolite, represented as *MMR*. By summing over these and normalizing to the concentration of the metabolite, we can find the required loss rate to balance the parameter choices for the model.

### Fitting

#### Sequential Monte Carlo - Approximate Bayesian Computing

Since the system is under-constrained, gradient based methods would find multiple solutions depending on the initial guess of the parameters. In order to explore the landscape of the parameters and determine if there is some underlying combinations more likely than others to give rise to the observed dynamics, Sequential Monte Carlo - Approximate Bayesian Computing [32] (SMC-ABC) was used to fit the system. In ABC the parameters are represented by a (posterior) distribution rather than a singular value, showing the probability (or likelihood) of the parameter being that value given the data. This approach allows for the inclusion of relevant information in the form of the prior, and gives a wider view the parameter space.

In the case of the GEEM model, the prior information takes the form of the ranges of observed values for the kinetics from literature. The distribution found can also indicate how sensitive a parameter may be, by the width and shape found. The basic algorithm is given in algorithm 1.

There are 2 main parts to the algorithm, the first being the initial generation of the valid candidate parameter arrays (henceforth called particle), and the second is resampling to cooldown to the posterior. In the initial particle generation, particles are sampled from the chosen prior distribution(s) in order to generate a test particle which contains the parameter values. These values are then run through the simulation, and the loss is calculated, using the summary statistic and the distance function, to evaluate the goodness of fit. If this loss is less than the threshold tolerance allowed at this step, the particle is accepted and added to the prior list. The prior is then resampled and this process is repeated until the desired number particles are generated.

In the resampling to cooldown to posterior step, the prior is first replaced with the new particles. Then the distribution kernel, which is a selected based on assumptions of the underlying distribution, is fit to the points. This distribution approximates the underlying distribution of the new samples and will be used to add noise based on the known distribution at this point. The tolerance is also updated to the next in the cooldown schedule. A sample is then randomly selected from the new prior and perturbed by noise generated by the distribution kernel. This becomes a new test particle and the model is simulated with these proposed parameters. The loss is calculated and if it is less than new tolerance, it is added to a new accepted particles list. The previous particles are again randomly sampled and noise from the distribution kernel is then added and the process is repeated. This continues until either the number of desired particles is found, or the compute resources are exhausted. If the desired number of particles is found, the tolerance is updated to the next in the cooldown schedule, the previous priors (particles) are replaced by the new points and the process is repeated until either the compute resources run out, or the desired tolerance is achieved. If the compute time is reached or the desired tolerance is reached, the found particles from the last completed step are returned.

##### Algorithm 1

Sequential Monte Carlo - Approximate Bayesian Computing

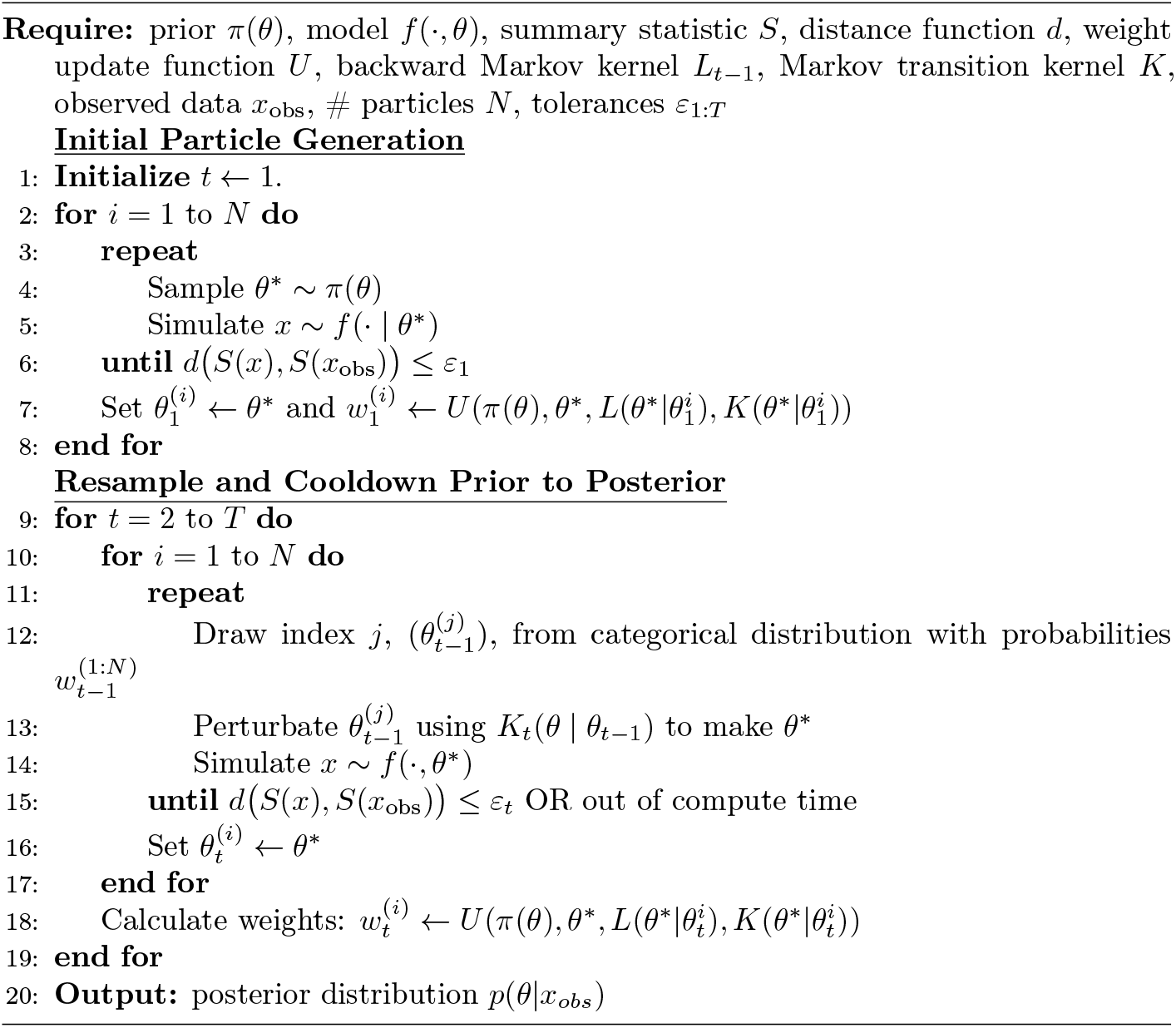

To perform the fitting, the prior, summary statistic, distance function, distribution kernel type, number of particles, cooldown schedule and compute resources need to be selected. In this experiment a uniform distribution was chosen for the prior distribution as it simplifies the calculations for the kernel and we do not have information to suggest a more characterized distribution. The summary statistic chosen must be able to summarize the results of the model in a way that can be evaluated with the distance function. Since we are evaluating our model against the experimental measurements, the summary statistic will be the model output at the time points of the experimental measurements. The distance function compares the summary statistic and the measurements. Since we are interested in how close the model can fit the measurements, we opt to use the RMSE between the predictions and the mean of the experimental measurements. The distribution kernel choice is informed by prior information, however, as there is no prior information to give an alternative, the default gaussian distribution was used. The final hyperparameters to choose are the number of particles, the tolerance cooldown schedule and the compute resources. Please note, the parameters are very system specific and must be tuned each time. Additionally, a finer cooldown schedule resolution can allow for a better fit, at the cost of more compute time. The cooldown schedule used here was: [, 0.25, 0.175, 0.125, 0.1, 0.09, 0.08, 0.07, 0.0675, 0.065, 0.0625, 0.06, 0.0575, 0.055, 0.0525, 0.05, 0.0475, 0.045, 0.0425, 0.04, 0.0375, 0.035, 0.0325]. The number of points can allow for more exploration but too many can cause the fitting process to miss rare areas of improvement. This can be overcome by increasing the compute resources, how many particles/iterations are tried before termination. In this experiment we settled on 1000 points as a compromise for efficiency and exploration, while the compute time was limited to 10000 attempts per thread before termination.

#### Enzyme Kinetics Fitting Parameters

In order to fit the GEEM model using SMC-ABC, a prior is required for the kinetic constants that are being fit. The priors for the enzyme synthesis and loss rates are consistent throughout the different versions of the GEEM model fit in this experiment. While no studies on the synthesis and loss rates for these specific enzymes were found, two studies were found which examined synthesis and loss rates in Arabidopsis. The enzyme synthesis rates reported by Piqques et al. [29] included various enzymes and their orders of magnitude were used as the priors. The enzyme loss rates reported by Li et al. [30] covered both specialized metabolite synthesis and stress response. These values formed the base for the corresponding priors. As the enzymes of interest were not present in the studies, the same priors were constructed for all enzymes across all models fit in this experiment, as shown in Table 2.

#### Metabolite Kinetics Fitting Parameters

The metabolic kinetic constants were handled differently through the versions of the experiment. The first version of the GEEM model used the ML predictions from Table 1 as the fixed values, referred further as ML predictions model.

To further improve the model, the next version of the GEEM model fit the kinetic constants using the ML predictions as priors, now referred to as the ML prior model. A range was created for each kinetic constant with priors set to one order of magnitude times the prediction, (*pred \** 10^1^) and below (*pred \** 10^*−*1^) allowing flexibility in the predictions, while maintaining the individual characteristics.

The final version of the model used the existing literature to further enhance the priors for the kinetic constants, referred to as Literature Enhanced Prior, by investigating the known kinetics of reactions involved in GSLs biosynthesis. As these reactions produce metabolites required under particular conditions (i.e. recruitment of specialized metabolites) it is assumed here that their reaction rates should be of a similar order of magnitude. Using enzymes with comparable functions, we identified the the minimum and the maximum *k*_*cat*_ were found and used these further widen the prior distributions.

The *K*_*M*_ ranges reported in literature ranged from 0.074*µM* [33] to 80000*µM* [22]. It is commonly assumed that the Michaelis constant should be on a scale similar to the concentration of the main substrate in the reaction. Textor et al. [34] reported that the *K*_*M*_ for the MAM1/3 reactions can be as high as 3000*µM*. To better align the typical orders of magnitude of the metabolites in our system, the *K*_*M*_ range was limited to the lower, more commonly observed value of 500*µM*. These priors were applied to all kinetics shown in Table 3.

## Results

The GEEM model was fit to the mock and JA treatments simultaneously for each of the three models versions described previously. Below we see the resulting metabolite response for each of the models as a function of the mRNA level for the mock and MeJA treatments. We first evaluate how well the machine learning model predicted kinetic constants were able to reproduce the data while fitting the enzyme kinetic parameters in isolation. Next, we made the model more flexible by also fitting the Michaelis Menten kinetic constants, using the ML predictions as a guide for the priors. Finally, we further widen the priors, by incorporating literature values of similar GSL reaction kinetics to further increase the models flexibility. Through progressively increasing the flexibility of the model, our aim was to create a model that can reproduce the experimentally observed data, while consistent with existing literature derived knowledge. In order to evaluate how each prior effects the models fit, the RMSE of the models found at the termination step of the SMC-ABC process was used, as well as a comparison of how the dynamics correspond to those observed in the data.

### Simulated Metabolite Results

#### ML Predictions

After predicting the kinetic constants using the machine learning model, the Michaelis Menten constants were fixed and the remaining enzyme kinetic parameters from the ODEs were fit to the data with a final mean RMSE of the models found of 0.0617. The resulting models reproduce the metabolite dynamics experimentally observed dynamics from both the mock and MeJA treatment (Fig 4). Comparing the simulations with the data, it can be seen that the approximately two-fold observed increase in 1MOI3M between the mock and the MeJA treatment was captured. The model results for 4MOI3M captures a similar increase between the mock and MeJA treatment observed in the data. However there is a slower response observed for the final metabolites (1MOI3M and 4MOI3M) in both conditions compared to experimental observations. Additionally, in the mock treatment, the simulated 1MOI3M concentrations are higher than those measured in the experimental data.

**Fig 4.**
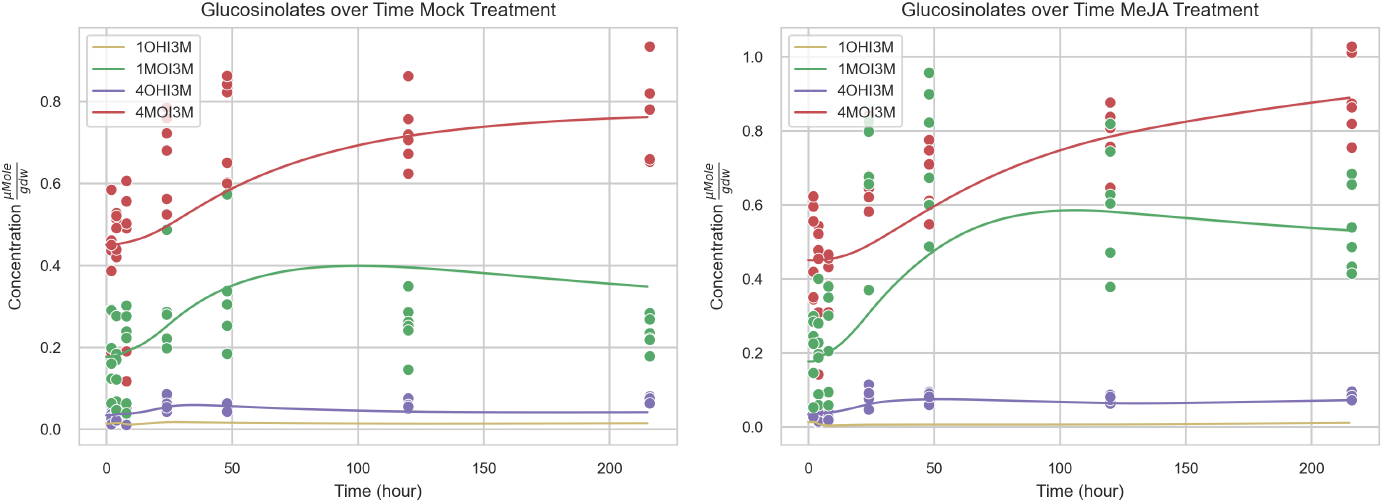
Metabolite content simulated by the ML predictions GEEM model. shows the model results of the mean glucosinolate concentrations over time for the mock (a/left) and MeJA (b/right) treatment with 95% CI shown (not visible due to high similarity to mean). The Michaelis Menten kinetics were fixed to the values predicted by the machine learning model. The mean RMSE of the models found was 0.0617.

#### ML Based Prior

Using the priors derived from the ML predictions, the models achieved an improved fit (mean RMSE of 0.0537) to the metabolite data (Fig 5). In particular, the post 24 h concentrations of 1MOI3M is better able to capture both treatments. The simulations of 4OHI3M and 4MOI3M also show a modest improvement in response compared to the models found by fixing the kinetic constants with ML predictions. However, the dynamics observed before 24 hours are still not well captured.

**Fig 5.**
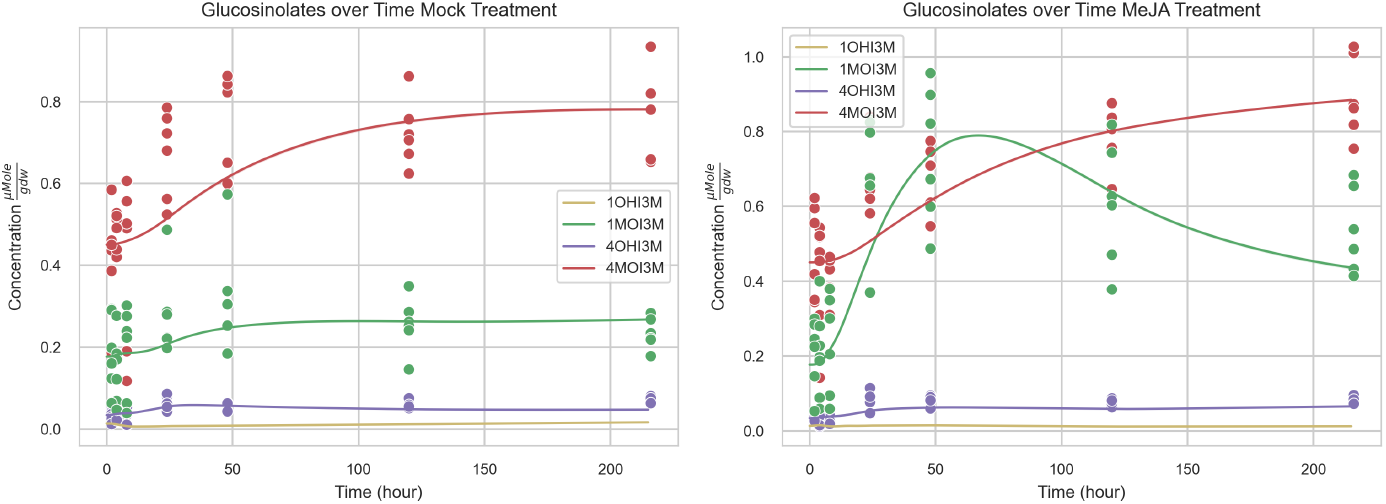
Metabolite content simulated by the ML priors GEEM model. shows the model results of the mean glucosinolate concentrations over time for the mock (a/left) and MeJA (b/right) treatment with 95% CI shown (not visible due to high similarity to mean). The Michaelis Menten kinetics were fit using priors derived from the machine learning model predictions. The mean RMSE of the models found was 0.0537.

#### Literature Enhanced Prior

After incorporation all available information, the machine learning model predictions and the literature observations, were combined into a broad prior for the metabolite kinetic predictions. This resulted in further improvements (mean RMSE of 0.0420). The metabolite simulation results (Fig 6) showed a further improvements to those obtained using the ML based prior model, including a higher concentration for 4OHI3M and lower 1MOI3M, concentrations which are closer to the actual data.

**Fig 6.**
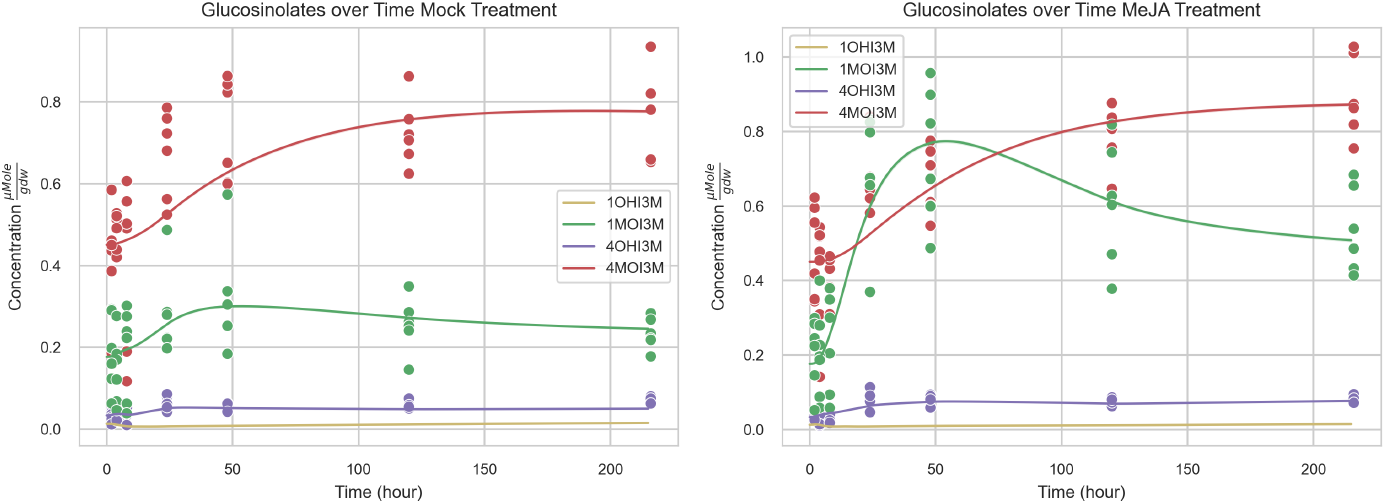
Metabolite content simulated by the Literature enhanced priors GEEM model. Figure shows the model results of the mean glucosinolate concentrations over time for the mock (a/left) and MeJA (b/right) treatment with 95% CI shown (not visible due to high similarity to mean). The Michaelis Menten kinetics were fit using priors derived from the combination of the machine learning model predictions and literature. The mean RMSE of the models found was 0.0420.

### Simulated Enzyme Results

In order to control the metabolite changes through time, and connect the mRNA levels to the metabolites, the enzyme concentrations are simulated. While there is no available experimental observations for the enzyme concentrations to validate against at this time, in order to asses the variability of the model, and examine the simulated dynamics, we show an example of the simulated enzyme concentrations from two of the models, the ML parameters (Fig 7) and the ML based priors (Fig 8) models.

**Fig 7.**
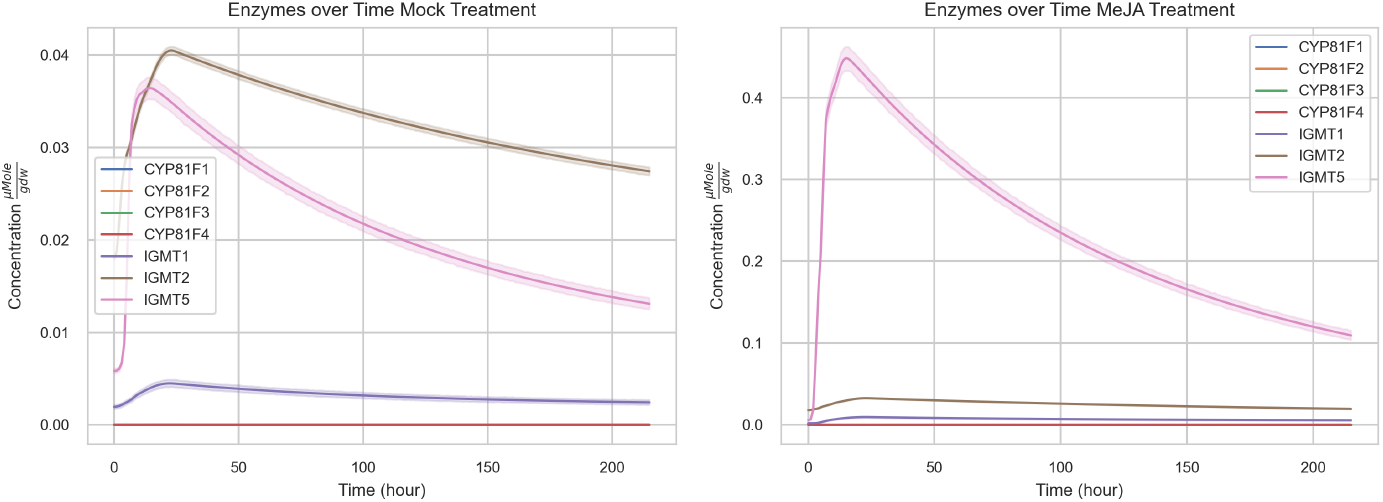
Enzyme content simulated by the fixed ML predictions GEEM model. shows the model results of the mean enzyme concentrations over time for the mock (a/left) and MeJA(b/right) treatment with 95% CI shown. The Michaelis Menten kinetics were fixed to the values predicted by the machine learning model.

**Fig 8.**
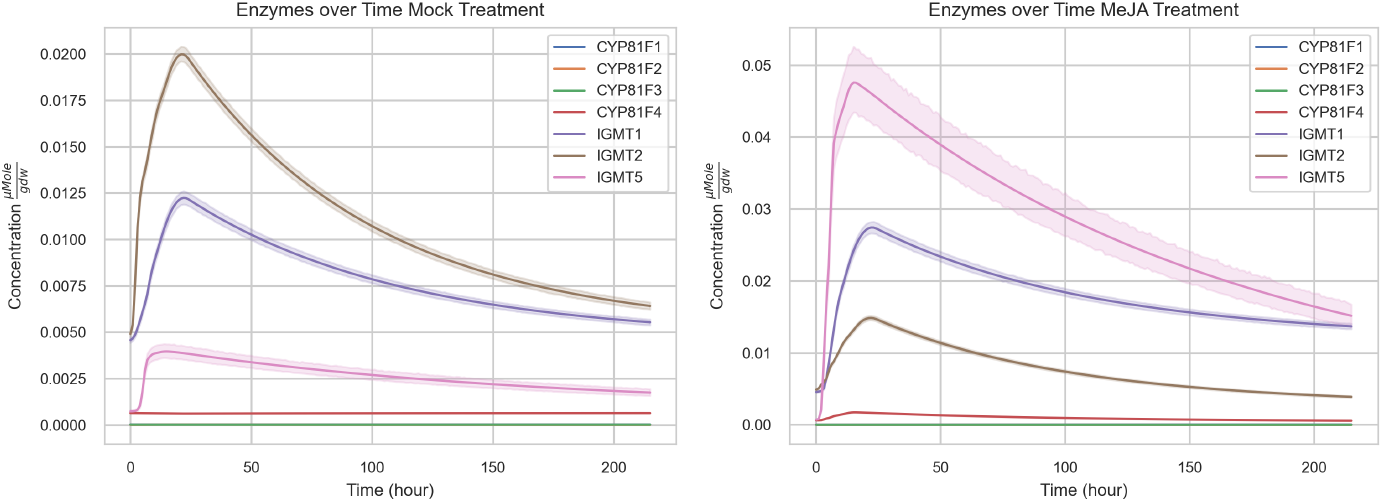
Enzyme content simulated by ML derived priors GEEM model. shows the model results of the mean enzyme concentrations over time for the mock (a/left) and MeJA (b/right) treatment with 95% CI shown. The Michaelis Menten kinetics were fit using priors derived from the machine learning model predictions.

The simulated enzyme concentrations exhibit different orders of magnitude across the two models. In the ML fixed model (Fig 7) the simulated enzymes have orders of magnitude difference between each other. In contrast, the ML priors model (Fig 8) shows the simulated enzyme concentrations for the mock treatment on a similar scale, but retaining the a larger difference in the MeJA treatment caused by the large difference in the mRNA level. In both models IGMT5 shows an approximate 10 fold increase between the mock and MeJA treatment as expected from the mRNA level data.

### Parameter Distributions

The resulting parameter distributions from the combined ranges models are shown in Fig 9. When comparing to the original bands seen in Table 3, the parameters have shifted to a more narrow distribution. While the orders of magnitude of the distributions vary between parameters, these figures show that during fitting the parameter space is refined to a narrower subset which can explain the experimental data dynamics.

**Fig 9.**
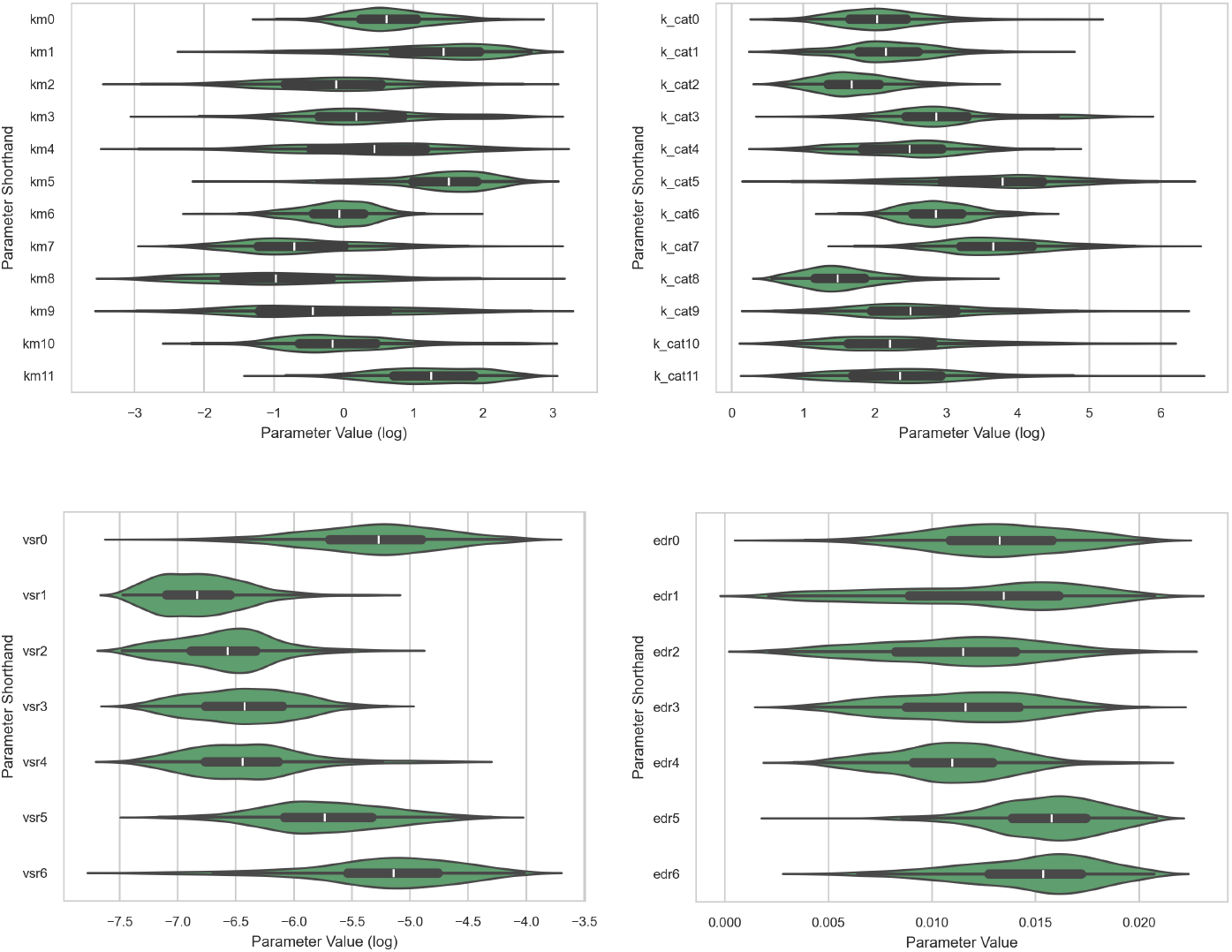
Parameter distributions example from the GEEM model. shows the parameter distributions found through the SMC-ABC fitting of the indolic GEEM model using the combined literature and ML parameters. The figures show for each reaction the distribution for that parameter type with the x axis being the parameter value in the scale of the prior, and the y axis being the shorthand names for the parameters. The figures show *k*_*M*_ (a/top left), *k*_*cat*_ (b/top right), enzyme synthesis rate (c/bottom left) and enzyme degredation rate (d/bottom right).

## Discussion

To test the hypothesis that changes in glucosinolate concentration can be explained by the observed changes in the mRNA levels of the biosyntehtic enzymes, a multilevel model was fit to de-novo experimental metabolite data of *A. thaliana* plants subjected to two treatments, over time. The models presented above all demonstrate the ability to reproduce the experimental metabolite dynamics. The quality of this fit is improved in each subsequent model by adding more information from existing literature. The fits indicate that a subset of the parameter space, rather than a single parameter vector, can reproduce the experimental observations.

This work extends models from Knoke et al. [10] by incorporating the time component, as well; as by including multiple conditions. While the model from Cloutier et al. [13] also utilizes an mRNA to metabolite model, we add to this model by including multiple conditions and by modeling the metabolites with Michaelis Menten dynamics, to attempt to find a measurable, physical constant. Additionally, we combine the model with current ML methods [15, 16] to try to take advantage of the new developments in ML. Our GEEM model shows that the inferred kinetics can reproduce metabolite dynamics under various conditions, such that it replicates the experimental dynamics. By including these different conditions, we further refined the kinetic constant predictions, and identified a subset of parameters that satisfy both the stressed and control (mock) conditions.

### Glucosinolate Content

As the flexibility of the model was increased by including more biologically grounded information, the quality of the fit to the metabolite data, and the flexibility of the system increased. The starting point of the GEEM model utilized the Michaelis Menten kinetic constants predicted using the machine learning model as fixed values. Fig 4 shows that while the ML predictions GEEM model can reproduce the overall experimental results, the MeJA treatment fit is missing much of the early dynamics. The figure also shows that the model overshoots the experimental data for 1MOI3M in the mock treatment.

Next the Michaelis Menten kinetic constants of the model were fit, using the machine learning model predictions as the guide for the priors for each of the kinetic constants, alongside the enzyme kinetics. This resulted in a better RMSE (Fig 5) as by fitting the Michaelis Menten kinetic constants, the ML priors model is better able to reproduce the early dynamics in both treatments. While 4OHI3M and 4MOI3M show similar, but improved, fits to the data, the concentration of 1MOI3M in both treatments shows a significant improvement. The mock treatment, 1MOI3M no longer overshoots the data, while the MeJA treatment captures the spike around 48 h.

Using the combination of the ML predictions and the literature-derived kinetics (Fig 6) achieves further improvements to the RMSE. The pre-50 h dynamics of 1MOI3M are better captured and the concentration of 4OHI3M better fits the experimental results. This indicates that while the ML based predictions contain some relevant information or prediction capabilities, more knowledge from literature still can improve the quality of the fit.

In all GEEM models fit, concentrations of pathway intermediates are lower than those observed in the mock treatment, likely due to the model constrained demand for specialized metabolites and the constraints on loss rates imposed by the initial conditions and parameter choices. The loss rate threshold also limits the dynamics as certain combinations yield values which fall outside of this range but would be a better fit. It is possible that the loss also changes due to the stressed condition, such as through more transporters or differences in degradation rates. A variable loss rate could help alleviate this problem but would require more information about the loss dynamics.

In the GEEM models fit, the early dynamics, pre 50 hour, show a slower rate of change than observed in the data. If we examine the metabolite data, there is a large variability in the measurements, making those initial values and early dynamics less clear, making the early dynamics more difficult to fit. If we consider the model, the differences could be from multiple sources. Glucosinolates are also produced in the roots, and also stored in other parts of the plant as well. The additional GSLs could originate in other parts and be transported to the leaves. We also limited the parameter priors from all values observed in the GSL pathway in literature, which could have excluded the real values. There could be missing enzymes involved in the process, or a co-factor or co-enzyme accelerating the reaction that is not modeled here. It is also possible that there are enzymes which are inhibited before the stressor, and then activated so the response is faster. As there is no information about these processes we elected to not include them in the model at this time. The model also uses the simplified form of Michaelis Menten kinetics, which also does not take into consideration competitive inhibition. In order to further refine this model, further experiments and analysis of the biosynthesis pathways should be conducted and the model updated with the additional information.

Together, these results show that the GEEM framework can mechanistically link transcriptional responses to time-resolved changes in indolic glucosinolate levels under stress. By clarifying how regulatory changes and kinetic constraints shape the accumulation of individual indolic glucosinolates over time, the GEEM model framework provides a clearer picture of stress-induced metabolic changes and can support the design of targeted experiments or breeding strategies to optimize these responses.

### Enzyme Content

The enzyme concentrations observed in the different fits can differ greatly in scale as seen in Figs 7 and 8. This is in part due to the low mRNA levels for some of the enzymes. Since the synthesis rate constant is derived from the initial mRNA level using the steady state assumption (see supplemental). If the mRNA level is very low, the synthesis rate will become very large.

The simulated enzyme concentrations (Figs 7 and 8) predict high concentrations of IGMT5 in the MeJA treatment due to the large increase in mRNA compared to the initial conditions, showing that the model is working as expected with the inputs received. After approximately 24 hours, the enzyme levels begin returned to their pre-treatment levels [23]. The higher concentrations of CYP81F4 and IGMT5 indicate that these are important to the stress response. This is also reflected in the mRNA levels, where the levels of these mRNA rise when exposed to MeJA. It can be seen that the Enzyme concentration of IGMT5 during MeJA treatment is an order of magnitude higher than the other enzymes. One reason the for this, in addition to the heightened response during MeJA treatment, is the low concentrations at the beginning of the experiment and the large increase in counts through the experiment. As the levels are low to begin with, the *k*_*sE*_ calculated for these enzymes are higher than the others, causing this large response.

In this model we make a quasi steady state assumption that the concentration of mRNA, enzymes, and metabolites are approximately at steady state before perturbation. While some of the genes have been shown to have some link to known circadian transcription factors, such as IGMT2/5 and CYP8F1/2 [31], it is not clear how strong these influences are and if they effect the concentration of the enzyme in a significant way. In order to further improve the model, these influences could be further examined and added into the model.

Due to a lack of proteomics data, it is difficult to validate the enzyme concentrations, hence the large difference in scale observed between the resulting fits seen in Figs 7 and This highlights that the models is still highly flexible due to the unknown dynamics of the proteins over the course of the experiment. While systems can be found which fit the metabolite data, as the enzyme concentrations is unconstrained, the Michaelis Menten kinetics parameters can vary greatly between similarly performing models (Fig 9). In order to achieve a more accurate model, it would be valuable to obtain initial enzyme concentration data in plants to impose stronger constraints on relative enzyme abundances. In addition, time-series measurements of enzyme levels and metabolites would allow further constraining of the model, improving its ability to capture enzyme dynamics and refining the metabolite predictions.

### Kinetic Constant Distributions

Examining at the distributions found from the literature enhance priors model in Fig 9, while some parameters appear to be approaching a more clustered distribution, many still have a wide distribution. This shows that with the constraints in the model currently, there are many different kinetic constants which could allow for the dynamics observed. Without more information on the kinetic constants or enzyme abundance to constrain the model, the reactions become semi-interchangeable. As previously mentioned the unconstrained enzyme concentrations allow for a wide range of possible kinetics, the two networks compensating in turn for the varied values.

The *K*_*M*_ values found via fitting (Fig 9a) are on a similar order of magnitude as the concentration of the substrate, as suggested by the hypothesis of enzymes co-evolving with the substrates and organism. From the literature it can be seen that the turnover rate (*k*_*cat*_) should be slower for membrane bound enzymes such as CYP81F1-4. Figs 9b shows that while the values found are on a similar scale to similar enzymes in the CYP family, they are not slower than the IGMT enzymes. The *K*_*M*_ of these CYP enzymes were also checked and appear to be on a similar scale, further supporting the model.

An advantage of using SMC-ABC is the incorporation of additional information through the priors. In this study we included the information as a uniform prior, with ranges derived from a combination of machine learning model predictions and literature. If later there is additional information uncovered which shows the prior could follow a certain distribution, this can be used improve the model.

If some of the kinetic constants were to be determined via kinetics experiments, the solution space would also reduced. This would exclude a large number of these models, allowing for a more targeted search of the space. The more kinetics given, the narrower the space would become. Additionally, if some of the concentrations for enzymes were known for nominal growth conditions, or if there was a time series of proteomics performed with the experimental data the model could be further improved. The nominal enzyme concentrations could be used to calibrate the initial conditions, linking the synthesis and degradation rates of the enzyme. If the full time-series was available, then the model could either be validated using this data, or it could be used as an additional metric to fit the model.

### Final Thoughts and Future Work

Our GEEM models shows that the observed changes in specialized metabolite concentrations can be modeled as a function of the mRNA counts under multiple conditions. By combining enzyme substrate kinetics predictions from machine learning models with literature knowledge, we are able to fit a model which can reproduce experimental observations of glucosinolate content in Arabidopsis using the mRNA levels as the model input. We have shown that while ML predictions alone can give a hint to the dynamics, by fitting the Michaelis Menten kinetics alongside the enzyme dynamics, the model can be further improved. Further augmentation of the priors by adding information from literature yields further improvements. By fitting across multiple conditions, we aim to improve the robustness of the inferred kinetics and better capture the dynamic behavior of the system, compared to previous works.

While the model can replicate the experimental observations, the distributions of the parameters found are still wide. While previous work has given a single solution [10], we show that given such an under-constrained system, many possible solutions can give rise to the observed dynamics. In order to further constrain the models, we propose including additional conditions in the fitting, which would give further constrains to the models. Additionally, if enzyme content data could be added to the fitting process, it would further help constrain and validate the enzyme concentrations of the models. Finally, in the GEEM model we do not consider competitive binding or feedback regulation in the Michaelis Menten kinetics. Adding this could yield a better representation of the system, but would require knowledge of binding preferences. We identified a number of possible sources for the slower early dynamics which can be investigated and used to improve the GEEM model.

The increasing availability of transcriptomic and metabolomic data highlights the need for methods that can effectively integrate these layers of information. By leveraging mathematical modeling to combine mRNA and metabolite measurements, we present a framework for capturing the dynamics of underlying metabolic networks. This approach is readily transferable to other systems and provides a foundation for more comprehensive analysis of multi-omics data.

## Acronyms

1MOI3M: 1-Methoxy-3-indolylmethyl Glucosinolate
1OHI3M: 1-hydroxyindole-3-ylmethyl Glucosinolate
4MOI3M: 4-Methoxy-3-indolylmethyl Glucosinolate
4OHI3M: 4-hydroxyindole-3-ylmethyl Glucosinolate
GSL: Glucosinolate
I3M: Indole-3-ylmethyl Glucosinolate
JA: Jasmonic Acid
MeJA: Methyl Jasmonate
ODE: Ordinary Differential Equation
SMC-ABC: Sequential Monte Carlo - Approximate Bayesian Computing

## Acknowledgments

Thanks goes to the lab technicians Jeremy Liu, Ringo van Wijk and Juliette Silven who supported the material and data collection. Additional thanks goes to Jordi Alonso Esteve, Julia Ruiz Capella, Samara Almeida Landman and the rest of the CropXR consortium.

This publication is part of the long-term program PlantXR: A new generation of breeding tools for extra-resilient crops (KICH3.LTP.20.005) which is financed by the Dutch Research Council (NWO), the Foundation for Food & Agriculture Research (FFAR), companies in the plant breeding and processing industry, and Dutch universities. These parties collaborate in the CropXR Institute (www.cropxr.org) that is funded through the National Growth Fund (NGF) of the Netherlands.

## Metabolite Data Collection

### Plant growth conditions and treatment

*Arabidopsis thaliana* ecotype Columbia-0 (Col-0) was used. Seeds were stratified in 0.1% agar at 4 °Cfor 48 h and sown on sand in closed trays with transparent lids to ensure 100% humidity under a 14-h long day. After 2 weeks, seedlings were transferred to individual pots containing a twice autoclaved soil: sand mixture (12:5). Plants were cultivated under a 10-hour long day at 21°*C*, 70% relative humidity and a light intensity of 200 *µmolm*^*−*2^*s*^*−*1^. Plants were watered three times per week and supplied with a nutrient solution once per week.

For treatment, rosettes of 5-week-old Arabidopsis were dipped for 3 sec in either a MeJA (0.1 mM; Merck Life Science BV) or a mock solution. Both solutions were prepared in tap water and contained 0.015% (v/v) Silwet L77 and 0.1% (v/v) ethanol. Samples were collected at 2, 4, 8, 24, 48, 120 and 216 h after treatment, as well as immediately after treatment (0 h) and 1 hour prior to treatment (-1 h) For each replicate one entire rosette was harvested and flash frozen in liquid nitrogen.

### Glucosinolate isolations and measurments

Sample were freeze dried overnight (Scanvac CoolSafe) and homogenized with a tissue lyser. From each sample, 15 mg of powdered tissue was transferred to a new tube and extracted with 0.4 mL of 100% UHPLC/MS-grade methanol (Biosolve BV, Valkenswaard, The Netherlands) containing 100pg/µL of the internal standards sinalbin (Phytoplan Diehm & Neuberger GmbH, Heidelberg, Germany) and sinigrin (Toronto Research Chemicals). Samples were vortexed thoroughly, followed by the addition of 0.8 mL HPLC-grade chloroform (Rathburn Chemicals Ltd., Walkerburn, Scotland) and further vortexing. The samples were incubated on ice for 10 min, after which 0.4 mL of ice-cold LC–MS-grade water (Biosolve BV, Valkenswaard, The Netherlands) was added. Samples were vortexed for 2 min and centrifuged at 600 rpm for 10 min at 4 °C.

The methanol:water phase was collectedby transferring 0.4 ml supernatant to a new tube, diluted with 1ml LCMS grade H2O and freeze dried overnight. Dried extracts were redissolved in 0.4 ml LCMS grade H2O, filtered trough a 0.22 µm microspin filter (BGB Analytik Vertrieb GmbH, Rheinfelden, Germany) and transferred to LCMS vails for analysis.

1ul per sample was run on U(H)PLC triple quad MS (1290 Infinity II, 6470 LC/TQ, Agilent) on a Poroshell 120 PFP column with a 2.1 × 5 mm 1.9 micron U(H)PLC guard column (Agilent). A program using acetonitrile (Biosolve BV, Valkenswaard, The Netherlands) and 0.1% formic acid in water (Biosolve BV, Valkenswaard, The Netherlands) was used. The program starts at 5% ACN followed by a gradient from 5% to 55% (v/v) acetonnitrile in 7,5 minutest, a gradient from 55% to 90% (v/v) acetonnitrile in 2 minutes and a gradient from 90% to 5% (v/v) acetonnitrile in 6 seconds. For quantification external standard dilutions were used of 4-hydroxyglucobrassicin, 4-methoxyglucobrassicin, neoglucobrassicin and glucobrassicin (Phytoplan Diehm & Neuberger GmbH, Heidelberg, Germany).

For the experiment, the concentrations of I3M, 1MOI3M, 4OHI3M and 4MOI3M were collected at 2, 4, 8, 24, 48, 120 and 216 hours after treatment, as well as immediately after treatment (0h) at 21°*C* with a mock and a meJA treatment as previously described. Six replicates were collected for each timepoint and each treatment which can be seen in Fig 2.

## References

1. Shakour ZT, Shehab NG, Gomaa AS, Wessjohann LA, Farag MA. Metabolic and Biotransformation Effects on Dietary Glucosinolates, Their Bioavailability, Catabolism and Biological Effects in Different Organisms. Biotechnology Advances. 2022 Jan;54:107784. doi:10.1016/j.biotechadv.2021.107784.

2. Hopkins RJ, Van Dam NM, Van Loon JJA. Role of Glucosinolates in Insect-Plant Relationships and Multitrophic Interactions. Annual Review of Entomology. 2009 Jan;54(1):57–83. doi:10.1146/annurev.ento.54.110807.090623.

3. Kissen R, Eberl F, Winge P, Uleberg E, Martinussen I, Bones AM. Effect of Growth Temperature on Glucosinolate Profiles in Arabidopsis Thaliana Accessions. Phytochemistry. 2016 Oct;130:106–18. doi:10.1016/j.phytochem.2016.06.003.

4. Justen VL, Fritz VA. Temperature-Induced Glucosinolate Accumulation Is Associated with Expression of BrMYB Transcription Factors. HortScience. 2013 Jan;48(1):47–52. doi:10.21273/HORTSCI.48.1.47.

5. Mewis I, Tokuhisa JG, Schultz JC, Appel HM, Ulrichs C, Gershenzon J. Gene Expression and Glucosinolate Accumulation in *Arabidopsis Thaliana* in Response to Generalist and Specialist Herbivores of Different Feeding Guilds and the Role of Defense Signaling Pathways. Phytochemistry. 2006 Nov;67(22):2450–62. doi:10.1016/j.phytochem.2006.09.004.

6. Brader G, Tas É, Palva ET. Jasmonate-Dependent Induction of Indole Glucosinolates in Arabidopsis by Culture Filtrates of the Nonspecific PathogenErwinia Carotovora. Plant Physiology. 2001 Jun;126(2):849–60. doi:10.1104/pp.126.2.849.

7. Kim JH, Jander G. Myzus Persicae (Green Peach Aphid) Feeding on Arabidopsis Induces the Formation of a Deterrent Indole Glucosinolate. The Plant Journal: For Cell and Molecular Biology. 2007 Mar;49(6):1008–19. doi:10.1111/j.1365-313X.2006.03019.x.

8. Erb M, Meldau S, Howe GA. Role of Phytohormones in Insect-Specific Plant Reactions. Trends in Plant Science. 2012 May;17(5):250–9. doi:10.1016/j.tplants.2012.01.003.

9. Tomczak JM, Węglarz-Tomczak E. Estimating Kinetic Constants in the Michaelis–Menten Model from One Enzymatic Assay Using Approximate Bayesian Computation. FEBS Letters. 2019 Oct;593(19):2742–50. doi:10.1002/1873-3468.13531.

10. Knoke B, Textor S, Gershenzon J, Schuster S. Mathematical Modelling of Aliphatic Glucosinolate Chain Length Distribution in Arabidopsis Thaliana Leaves. Phytochemistry Reviews. 2009 Jan;8(1):39–51. doi:10.1007/s11101-008-9107-3.

11. Cloutier M, Chen J, Tatge F, McMurray-Beaulieu V, Perrier M, Jolicoeur M. Kinetic Metabolic Modelling for the Control of Plant Cells Cytoplasmic Phosphate. Journal of Theoretical Biology. 2009 Jul;259(1):118–31. doi:10.1016/j.jtbi.2009.02.022.

12. Pagare S, Bhatia M, Tripathi N, Pagare S, Bansal Y. Secondary Metabolites of Plants and Their Role: Overview. Current Trends in Biotechnology and Pharmacy. 2015:293–304.

13. Cloutier M, Xiang D, Gao P, Kochian LV, Zou J, Datla R, et al. Integrative Modeling of Gene Expression and Metabolic Networks of Arabidopsis Embryos for Identification of Seed Oil Causal Genes. Frontiers in Plant Science. 2021 Apr;12:642938. doi:10.3389/fpls.2021.642938.

14. Hickman R, Van Verk MC, Van Dijken AJH, Mendes MP, Vroegop-Vos IA, Caarls L, et al. Architecture and Dynamics of the Jasmonic Acid Gene Regulatory Network. The Plant Cell. 2017 Sep;29(9):2086–105. doi:10.1105/tpc.16.00958.

15. Kroll A, Engqvist MKM, Heckmann D, Lercher MJ. Deep Learning Allows Genome-Scale Prediction of Michaelis Constants from Structural Features. PLOS Biology. 2021 Oct;19(10):e3001402. doi:10.1371/journal.pbio.3001402.

16. Kroll A, Rousset Y, Hu XP, Liebrand NA, Lercher MJ. Turnover Number Predictions for Kinetically Uncharacterized Enzymes Using Machine and Deep Learning. Nature Communications. 2023 Jul;14(1):4139. doi:10.1038/s41467-023-39840-4.

17. Pfalz M, Mikkelsen MD, Bednarek P, Olsen CE, Halkier BA, Kroymann J. Metabolic Engineering in Nicotiana Benthamiana Reveals Key Enzyme Functions in Arabidopsis Indole Glucosinolate Modification. The Plant Cell. 2011 Feb;23(2):716–29. doi:10.1105/tpc.110.081711.

18. Pfalz M, Mukhaimar M, Perreau F, Kirk J, Hansen CIC, Olsen CE, et al. Methyl Transfer in Glucosinolate Biosynthesis Mediated by Indole Glucosinolate O-Methyltransferase 5. Plant Physiology. 2016 Dec;172(4):2190–203. doi:10.1104/pp.16.01402.

19. Kroll A, Lercher MJ. DLKcat Cannot Predict Meaningful Kcat Values for Mutants and Unfamiliar Enzymes. Biology Methods and Protocols. 2024 Jan;9(1):bpae061. doi:10.1093/biomethods/bpae061.

20. Rives A, Meier J, Sercu T, Goyal S, Lin Z, Liu J, et al. Biological structure and function emerge from scaling unsupervised learning to 250 million protein sequences. Proceedings of the National Academy of Sciences. 2021;118(15):e2016239118. BioRxiv 10.1101/622803. Available from: https://www.pnas.org/doi/full/10.1073/pnas.2016239118. doi:10.1073/pnas.2016239118.

21. Landrum G. RDKit: Open-Source Cheminformatics Software; 2016. Available from: https://github.com/rdkit/rdkit/releases/tag/Release_2016_09_4.

22. Andersson D, Chakrabarty R, Bejai S, Zhang J, Rask L, Meijer J. Myrosinases from Root and Leaves of Arabidopsis Thaliana Have Different Catalytic Properties. Phytochemistry. 2009;70(11-12):1345–54. doi:10.1016/j.phytochem.2009.07.036.

23. Sasaki Y, Asamizu E, Shibata D, Nakamura Y, Kaneko T, Awai K, et al. Monitoring of Methyl Jasmonate-Responsive Genes in Arabidopsis by cDNA Macroarray: Self-Activation of Jasmonic Acid Biosynthesis and Crosstalk with Other Phytohormone Signaling Pathways. DNA research: an international journal for rapid publication of reports on genes and genomes. 2001 Aug;8(4):153–61. doi:10.1093/dnares/8.4.153.

24. Islam S, Kjällquist U, Moliner A, Zajac P, Fan JB, Lönnerberg P, et al. Characterization of the Single-Cell Transcriptional Landscape by Highly Multiplex RNA-seq. Genome Research. 2011 Jul;21(7):1160–7. doi:10.1101/gr.110882.110.

25. Hashimshony T, Wagner F, Sher N, Yanai I. CEL-Seq: Single-Cell RNA-Seq by Multiplexed Linear Amplification. Cell Reports. 2012 Sep;2(3):666–73. doi:10.1016/j.celrep.2012.08.003.

26. Takahashi M, Morikawa H. Nitrogen Dioxide at Ambient Concentrations Induces Nitration and Degradation of PYR/PYL/RCAR Receptors to Stimulate Plant Growth: A Hypothetical Model. Plants. 2019 Jul;8(7):198. doi:10.3390/plants8070198.

27. Tolleter D, Smith EN, Dupont-Thibert C, Uwizeye C, Vile D, Gloaguen P, et al. The Arabidopsis Leaf Quantitative Atlas: A Cellular and Subcellular Mapping through Unified Data Integration. Quantitative Plant Biology. 2024 Feb;5:e2. doi:10.1017/qpb.2024.1.

28. Wuyts N, Massonnet C, Dauzat M, Granier C. Structural Assessment of the Impact of Environmental Constraints on Arabidopsis Thaliana Leaf Growth: A 3D Approach. Plant, Cell & Environment. 2012;35(9):1631–46. doi:10.1111/j.1365-3040.2012.02514.x.

29. Piques M, Schulze WX, Höhne M, Usadel B, Gibon Y, Rohwer J, et al. Ribosome and Transcript Copy Numbers, Polysome Occupancy and Enzyme Dynamics in Arabidopsis. Molecular Systems Biology. 2009 Oct. doi:10.1038/msb.2009.68.

30. Li L, Nelson CJ, Trösch J, Castleden I, Huang S, Millar AH. Protein Degradation Rate in Arabidopsis Thaliana Leaf Growth and Development. The Plant Cell. 2017 Feb;29(2):207–28. doi:10.1105/tpc.16.00768.

31. Romero-Campero FJ, Pedro, Romero-Losada AB. ATTRACTOR: Fiesta release of ATTRACTOR. Zenodo; 2020. Available from: https://doi.org/10.5281/zenodo.3780022. doi:10.5281/zenodo.3780022.

32. Sisson SA, Fan Y, Tanaka MM. Sequential Monte Carlo without Likelihoods. Proceedings of the National Academy of Sciences. 2007 Feb;104(6):1760–5. doi:10.1073/pnas.0607208104.

33. Yin R, Frey M, Gierl A, Glawischnig E. Plants Contain Two Distinct Classes of Functional Tryptophan Synthase Beta Proteins. Phytochemistry. 2010 Oct;71(14):1667–72. doi:10.1016/j.phytochem.2010.07.006.

34. Textor S, Bartram S, Kroymann J, Falk KL, Hick A, Pickett JA, et al. Biosynthesis of Methionine-Derived Glucosinolates in Arabidopsis Thaliana : Recombinant Expression and Characterization of Methylthioalkylmalate Synthase, the Condensing Enzyme of the Chain-Elongation Cycle. Planta. 2004 Apr;218(6):1026–35. doi:10.1007/s00425-003-1184-3.

35. Textor S, de Kraker JW, Hause B, Gershenzon J, Tokuhisa JG. MAM3 Catalyzes the Formation of All Aliphatic Glucosinolate Chain Lengths in Arabidopsis. Plant Physiology. 2007 May;144(1):60–71. doi:10.1104/pp.106.091579.

36. Klein M, Reichelt M, Gershenzon J, Papenbrock J. The Three Desulfoglucosinolate Sulfotransferase Proteins in Arabidopsis Have Different Substrate Specificities and Are Differentially Expressed. The FEBS Journal. 2006;273(1):122–36. doi:10.1111/j.1742-4658.2005.05048.x.

